# APP binds to the EGFR ligands HB-EGF and EGF, acting synergistically with EGF to promote ERK signaling and neuritogenesis

**DOI:** 10.1101/2020.06.12.149062

**Authors:** Joana F. da Rocha, Luísa Bastos, Sara C. Domingues, Ana R. Bento, Uwe Konietzko, Odete A. B. da Cruz e Silva, Sandra I. Vieira

## Abstract

The amyloid precursor protein (APP) is a transmembrane glycoprotein central to Alzheimer’s disease (AD) with functions in brain development and plasticity, including in neurogenesis and neurite outgrowth. Epidermal growth factor (EGF) and heparin-binding EGF-like growth factor (HB-EGF) are well described neurotrophic and neuromodulator EGFR ligands, both implicated in neurological disorders like Schizophrenia and AD. Here we show that APP interacts with these two EGFR ligands and characterize the effects of APP-EGF interaction in ERK activation and neuritogenesis. HB-EGF was identified as a novel APP interactor in a yeast two-hybrid screen of a human brain cDNA library. Yeast co-transformation and co-immunoprecipitation assays confirmed APP interaction with HB-EGF. Moreover, co-immunoprecipitation also revealed that APP binds to cellular pro-EGF. Overexpression of HB-EGF in HeLa cells, or exposure of SH-SY5Y cells to EGF, both resulted in increased APP protein levels. EGF and APP were also observed to synergistically activate the ERK signaling pathway, crucial for early neuronal differentiation. Immunofluorescence analysis of cellular neuritogenesis in conditions of APP overexpression and EGF exposure, confirmed a synergistic effect in promoting the number and the mean length of neurite-like processes per cell. Synergistic ERK activation and neuritogenic effects were completely blocked by the EGFR inhibitor PD 168393, implying EGF-induced activation of EGFR as part of the mechanism. This work shows novel APP protein interactors and provides a major insight into the APP-driven mechanisms underlying neurite outgrowth and neuronal differentiation, with potential relevance for AD and for adult neuroregeneration.

## Introduction

The amyloid precursor protein (APP) is a type 1 integral membrane glycoprotein with a large extracellular domain and a short cytoplasmic region. The extracellular domain is organized in two N-terminal folded domains, the E1 and E2 domains, both containing a growth factor-like domain (GFLD) with heparin binding properties [1]. During its dynamic intracellular trafficking, APP undergoes proteolytic cleavages to release biologically active peptides [2]. Close to its membrane domain, APP can be cleaved by β-secretase or α-secretase, releasing soluble N-terminal fragments. The resulting membrane-anchored C-terminal fragments are subsequently cleaved by the γ-secretase complex to release two peptides: Abeta and the APP intracellular domain AICD, after β-secretase, or p3 and AICD, after α-secretase [3]. APP is a key protein in Alzheimer’s disease (AD), the most prevalent neurodegenerative disorder [4], and has been further implicated in several other neurological disorders and brain injury [5]. Highly enriched in neurons, APP appears to be critical for normal brain development and adult brain plasticity [6]. Specific roles in neurogenesis [7], neuronal migration [8], neurite outgrowth [9] and guidance [10], synaptogenesis [11], synaptic plasticity [12] and neuroregeneration [13, 14] have all been described.

Similar to APP, the pro-epidermal growth factor (pro-EGF) is a type I transmembrane glycoprotein cleaved by cell surface proteases to release the mature epidermal growth factor (EGF). The human EGF precursor has a large extracellular domain with 9 EGF-like domains, a helical transmembrane domain, and a shorter cytoplasmic domain [15, 16]. Pro-EGF is the classic member of the EGF family of growth factors that includes several members with highly similar domains, namely the EGF-like domain containing six cysteine residues [17]. One of the proteins included in this group is the pro-heparin-binding EGF-like growth factor (pro-HB-EGF), also a type I transmembrane glycoprotein cleaved to yield the soluble ligand HB-EGF. The C-terminal domain of pro-HB-EGF shares 40% sequence identity with pro-EGF, while the mature HB-EGF domain is described to be an N-terminally extended form of the mature EGF [18]. HB-EGF has a single EGF-like domain which partially overlaps with a heparin-binding domain, and a diphtheria toxin binding domain; these last two are distinct features of HB-EGF, not present in EGF [17, 18]. Importantly, both EGF and HB-EGF soluble and membrane-bound forms (their precursor proteins) have been described as biologically active, with membrane-bound forms competing with mature proteins for receptor binding [19]. Both EGF and HB-EGF bind directly to the epidermal growth factor receptor (EGFR; also known as ERBB1 or HER1), and HB-EGF can additionally bind ERBB4 (or HER4) [20]. Binding of the mature growth factor to EGFR induces receptor dimerization and auto-phosphorylation of the receptor tyrosine residues, which in turn recruits phosphotyrosine binding proteins and initiates different signaling events. The major pathways activated by EGFR include Ras stimulation of the RAF-MEK-ERK1/2 cascade, PI3K stimulation of Akt and PLC-gamma, and JAK stimulation of the STAT pathway [21]. EGF and HB-EGF are well described mitogens for multiple cell lines, ranging from epithelial cells and fibroblasts to neural stem cells [22, 23]. Given their neurotrophic and neuromodulatory roles, these molecules also promote cell differentiation, survival and plasticity [24–28], and have been implicated in neurological disorders like Schizophrenia, Parkinson’s and AD [23, 29–31].

During the years, different studies have punctually revealed an interplay between APP and EGF. EGF stimulation has been implicated in the regulation of APP processing. On one hand, EGF stimulation promotes sAPP release (including sAPPα) in rat PC12 cells, human epidermoid carcinoma cells, and mice adult subventricular zone (SVZ) progenitors [32–34]. On the other hand, EGF treatment was shown to promote β-secretase activity and increase Abeta production in rat and human immortalized cell lines expressing the familial AD-related APP Swedish mutant [35]. Neuronal cultures from transgenic animals overproducing the human APP carrying the Swedish mutation display alterations in EGFR degradation after EGF stimulation [36]. The AICD fragment was also reported to negatively regulate EGFR gene expression [37]. Interestingly, sAPPα is able to bind type-C cells of the adult SVZ that are actively dividing and express EGFR. While *in vivo* administration of sAPPα increased proliferation of these adult progenitors, anti-APP antibodies blocked their EGF-induced *in vitro* proliferation [32].

In this work, we present HB-EGF as a novel APP protein interactor, first identified in a yeast two-hybrid screen using a human cDNA library. This interaction was validated in a mammalian cell culture and its effects on APP protein expression analyzed. Due to EGF and HB-EGF partial overlapping in terms of cell/tissue expression and functions, a physical interaction between APP and pro-EGF was additionally tested and demonstrated. APP and EGF were observed to induce a synergistic effect on neuritogenesis via ERK1/2 activation, known to be required for neuronal differentiation and regeneration.

## Materials and Methods

### Antibodies and reagents

The following primary antibodies were used: rabbit anti-HB-EGF (antibodies-online), mouse anti-Myc (9B11; Cell Signaling Technologies), rabbit anti-EGF (antibodies-online), rabbit anti-pro-EGF (against aa 700-800; Novus Biologicals, R&D Systems Europe Ltd, England), rabbit anti-APP C-Terminal (clone CT695; Invitrogen), mouse anti-APP N-terminal (clone 22C11; Chemicon), mouse anti-APP residues 1-16 of Abeta (clone 6E10), rabbit anti-ERK1/2 (Millipore, Merck Life Science S.L.U., Portugal), rabbit anti-phosphorylated ERK1/2 (Thr202/Tyr204 Erk1 and Thr185/Tyr187 Erk2; Millipore), anti-acetylated alpha-tubulin (Sigma-Aldrich, Química S.A., Portugal), mouse anti-βIII Tubulin (Sigma-Aldrich). HRP-conjugated anti-mouse and anti-rabbit secondary antibodies were from GE-Healthcare. Secondary antibodies Alexa Fluor 405 goat anti-rabbit, Alexa Fluor 594 goat anti-rabbit or anti-mouse, and Alexa Fluor 488 goat anti-mouse were from Life Technologies. Human recombinant EGF was from Cell Signaling and was used at a 100 ng/mL working concentration (WC). The EGFR inhibitor drug PD 168393 (Sigma-Aldrich) was used at 10 μM WC. Texas Red-conjugated Transferrin was from Molecular Probes.

### Plasmid construction

The vector used to insert the bait cDNAs in the yeast two-hybrid (YTH) screens was Gal4 DNA binding domain (Gal4-BD) expression vector pAS2-1 (Clontech, Enzifarma, Portugal). Bait-1 cDNA, coding for full-length human APP (neuronal isoform 695), and bait-2 coding for the intracellular domain of human APP695 (AICD), were PCR amplified and inserted in the vector pAS2-1 using *NcoI/SmaI* restriction enzyme sites, in frame with the Gal4-BD. The HB-EGF cDNA was obtained from the Human Brain Matchmaker cDNA library (Clontech). Briefly, the pACT2-HB-EGF DNA was extracted from the positive clones obtained in the large-scale yeast mating [38, 39] and was rescued by transformation in *E. coli* XL-1 Blue. This HB-EGF cDNA was used to prepare the Myc-HB-EGF construct, by digestion with *Eco*RI and *Xho*I (New England Biolabs, NEB) and insertion into pCMV-Myc (Clontech), in frame with the Myc coding sequence. The human APP isoform 695 cDNA had been previously fused in frame with GFP in the pEGFP-N1 mammalian expression vector [40].

### Validation of APP-HB-EGF interaction by co-transformation in yeast

The interaction between APP and HB-EGF (and AICD-HBEGF) was verified by co-transformation of the bait (pAS2-1-APP or pAS2-1-AICD) and the prey plasmid (HB-EGF-pACT2) in the yeast strain AH109 (Clontech). First, the transformants were assayed for HIS3, ADE2, and MEL1 reporter genes’ intrinsic activation. Then yeast strain AH109 was co-transformed using the lithium acetate method, with the following plasmid pairs: pAS2-1/pACT2 empty vectors (negative control); pVA3-1/pTD1-1, expressing the Gal4-BD-p53 and the Gal4-AD-SV40 large T antigen fusions, respectively (positive control); pAS2-1/HB-EGF-pACT2; APP-pAS2-1/pACT2; AICD-pAS2-1/pACT2; APP-pAS2-1/HB-EGF-pACT2; AICD-pAS2-1/HB-EGF-pACT2. To confirm protein-protein interactions (reporter genes activation), fresh colonies of the co-transformants were assayed for growth on triple drop-out synthetic medium for yeast cells lacking Trp, Leu and His (SD/TDO) and quadruple drop-out synthetic medium for yeast cells lacking Trp, Leu, His and Ade (SD/QDO) plates, and later on SD/QDO containing 5-bromo-4-chloro-3-indolyl-α-d-galactopyranoside (X-α-Gal).

### Mammalian cell lines culture

The HeLa cell line was maintained in 10% heat inactivated fetal bovine serum (FBS, Gibco, Thermo Fisher Scientific) Dulbeco’s Minimal Essential Media (DMEM, Gibco), and the SH-SY5Y human neuroblastoma cell line was maintained in 10% FBS Minimum Essential/F12 medium (MEM/F12, Gibco). Both cells’ media included 1% antibiotic/antimycotic mix (Gibco), and cell cultures were kept in a humidified, 37°C, 5% CO_2_ incubator. For biochemical studies cells were plated in order to reach 60-80% confluency next day. For morphometric and immunocytochemistry studies cells were plated at an initial 2.0×10^4^ cells/cm^2^ density, onto six-wells plates containing sterile coverslips, and allowed to grow for 24h before experiments.

### Mammalian cells transfection

Mammalian cell lines were transfected with the TurboFect™ reagent according to the manufacturer’s instructions (Fermentas Life Sciences). The cDNA(s) were diluted in serum-free growth medium, and the cDNA/TurboFect™ ratio used was 1:2. The total amount of each reagent was adjusted accordingly to the monolayer of cells area. For co-transfection experiments 1/2 of each cDNA was used. The cDNA/TurboFect™ mixture was vortexed and incubated for 15-20 min at room temperature (RT), and then added dropwise to each well, with gentle rocking. Cells were further incubated for 6h at 37°C in a CO2 incubator, after which the medium was changed. Empty pEGFP-N1 and/or pCMV-Myc expression vector(s) (here named ‘V’ or ‘Vs’) were used as controls. Experimental conditions for the HB-EGF assays included single or co-transfection of human APP-GFP and Myc-HB-EGF, and the Myc and GFP empty vectors: 1) pEGFP and pCMV-Myc empty vectors (‘Vs’ condition), 2) Myc-HB-EGF cDNA and the pEGFP empty vector (‘Myc-HB-EGF’ condition), 3) APP-GFP cDNA and the pCMV-Myc empty vectors (‘APP-GFP’ condition), and 4) APP-GFP and Myc-HB-EGF cDNAs (‘APP-GFP/Myc-HB-EGF’ condition). Experimental conditions for the EGF assays included single transfections of human APP-GFP (‘APP-GFP’ or ‘APP’) and the GFP empty vector (‘V’).

### GFP-Trap co-precipitation assay

APP-GFP protein, overexpressed by HeLa cells following transient transfection, and their binding partners, were co-precipitated from cells extracts by pulling-down the GFP moiety with GFP-Trap^®^ (Chromotek), according to the manufacturer’s instructions. Briefly, after 24h of transfection with the APP-GFP cDNA, alone or co-transfected with the Myc-HB-EGF cDNA, cells were incubated with the DSP crosslinker (Pierce) for 45 min at RT. Cells were subsequently washed in phosphate buffer saline (PBS), and collected with a scrapper in 1 mL ice-cold phenylmethylsulfonyl fluoride (PMSF)-containing PBS, on ice. After a centrifugation step, the cell pellet was lysed for 30 min on ice with non-denaturant lysis buffer 1 (10 mM Tris-HCl pH 7.5, 0.5 mM EDTA, 0.5% Gepac-ca-630, 150 mM NaCl, 1 mM PMSF) supplemented with protease inhibitors cocktail (Sigma-Aldrich), 1 mM NaF, and 10 mM sodium orthovanadate. Following centrifugation (5 min at 20000 ×g, 4°C), the supernatant was collected, an aliquot of 25 μL was taken (‘cells lysates’ fraction), and total protein quantified (Pierce’s BCA protein kit; Thermo Scientific, Thermo Fisher Scientific). Mass-normalized lysates were pull-downed with GFP-Trap for 3h with orbital shaking at 4°C. The beads were magnetically separated until the supernatant was clear, and resuspended in wash buffer (10 mM Tris-HCl pH 7.5; 150 mM NaCl; 0.5 mM EDTA) for 10 min with agitation at 4°C. Magnetic separation and washing step were repeated 4 times, after which the proteins were eluted in Laemmli sample buffer.

### Cortex dissection and APP-pro-EGF co-Immunoprecipitation

Wistar Hannover rats (9-12 weeks) were obtained from Harlan Interfaune Ibérica, SL. The number of animals used and the suffering was minimized, and all the experiments met the European legislation for animal experimentation (2010/63/EU; European Council Directive 86/609/EEC). The experiment was approved and supervised by our Institutional Animal Care and Use Committee (IACUC): Comissão Responsável pela Experimentação e Bem-Estar Animal (CREBEA). Briefly, animals were sacrificed by cervical stretching followed by decapitation and their cortices dissected on ice. Tissues were further weighed and homogenized on ice with a Potter-Elvehjem tissue homogenizer in non-denaturating lysis buffer (50 mM Tris-HCl pH 8.0, 120 mM NaCL and 4% CHAPS) containing protease inhibitors [41]. The tissue extracts were used for immunoprecipitation analysis as described below (n = 2). Mice cortex dissection was performed as described above for rat, but tissue was homogenized in NP-40 based lysis buffer (n = 1).

For immunoprecipitation of rodent cortices, Dynabeads Protein G (Dynal, Invitrogen, Thermo Fisher Scientific) were washed in 3% bovine serum albumin (BSA) in PBS, and primary antibodies (rabbit anti-pro-EGF or mouse anti-APP 22C11) were incubated with Dynabeads according to the manufacturer’s instructions. Cortex lysates were precleared with 0.6 mg Dynabeads for 1h at 4°C with agitation and incubated with antibody-Dynabeads overnight (O/N) at 4°C with agitation. The immunoprecipitates were washed in PBS and proteins denatured by boiling in Laemmli loading buffer followed by SDS-PAGE and immunoblotting.

### SDS-PAGE and Western blot

The synergistic effect of EGF and APP in ERK1/2 activation was first studied in HeLa and SH-SY5Y cells transfected for 24h with the GFP vector (‘V’) or APP-GFP and exposed in the last 30 min to EGF. Cellular lysates were subjected to Western blot (WB) analysis as follows. Cells were washed and harvested with 1% boiling sodium dodecyl sulfate (SDS). SDS harvested cell lysates were further boiled for 10 min. Cell lysates were sonicated for 30 sec, and total protein content was measured as described above. When required, the conditioned medium was collected, cleared by centrifugation, and harvested into final 1% SDS samples. Aliquots for WB were mass-normalized to the total protein content of the respective cellular lysates (Pierce’s BCA kit; Thermo Scientific). Mass-normalized aliquots were subjected to electrophoresis on 5-20% gradient SDS-PAGE, and subsequently electrotransferred into nitrocellulose membranes. These were initially processed for Ponceau-S staining in order to assess gel loading [42], since in cell differentiation studies beta-actin and other cytoskeleton markers are not suitable loading controls [42–44]. Subsequently, primary antibodies were incubated as specified by the manufacturer’s instructions (from 2h to O/N) and secondary antibodies for 2h. Membranes were further washed 3× with TBS-T or PBS-T for 10 min, and submitted to the ECL^™^ (enhanced chemiluminescence; GE Healthcare) detection method, using X-ray films (GE Healthcare), or directly imaged by ChemiDoc™ touch Imaging System (Bio-Rad Laboratories, Lda, Amadora, Portugal).

Ponceau-S stained membranes and ECL X-ray films were scanned (GS-800 imaging densitometer, Bio-Rad) and protein immunoreactive bands quantified (Quantity One densitometry software or Image Lab software from Bio-Rad). WB data is presented as fold-increases of a control condition (stated in figure caption), corrected to Ponceau-S, used as loading control.

### RNA extraction, cDNA synthesis and quantitative real-time polymerase chain reaction

Total RNA was isolated from HeLa cells using NZYol reagent (NZYTech, Lisboa, Portugal). RNA concentration was determined using the DS-11 Series Spectrophotometer / Fluorometer (DeNovix, Wilmington, Delaware, USA). cDNA synthesis was performed using 1 μg of total RNA, RevertAid reverse transcriptase (Thermo Fisher Scientific, Waltham, Massachusetts, USA), and oligo-dT primers. Real-time polymerase chain reaction was carried out in duplicate using 2× SYBR^®^ Green Supermix (Bimake, Houston, Texas, USA), and reactions were run on a CFX Connect Real-Time PCR Detection System (BioRad Laboratories, Lda, Amadora, Portugal). Primer sequences were designed using Beacon Designer™ 7 (Premier Biosoft, Palo Alto, California, USA) for the APP and HPRT1 human genes. The HPRT1 mRNA levels were used as an internal control to normalize the mRNA levels of APP gene. The oligonucleotides used were: APP, forward 5’-CAATGTGGATTCTGCTGAT-3’ and reverse 5’-CCTCTGCTACTTCTACTACTT-3’; and HPRT1, forward 5’-GGCGTCGTGATTAGTGATG-3’ and reverse 5’-CAGAGGGCTACAATGTGATG-3’. Amplification efficiency for each primer pair was determined using serial dilutions of cDNA samples. No template controls (NTCs) were included in all experiments as negative controls. For gene expression analysis, 2 μL of 1:10 diluted cDNA was added to 10 μL of 2× SYBR^®^ Green Supermix (Bimake, Houston, Texas, USA) and the final concentration of each primer was 250 nM in 20 μL of total volume. The thermocycling reaction was initiated by activation of Taq DNA Polymerase by heating at 95°C during 3 min, followed by 40 cycles of a 12 s denaturation step at 95°C and a 30 s annealing/elongation step at 60°C. After the thermocycling reaction, the melting step was performed with slow heating, starting at 60°C and with a rate of 1%, up to 95°C, with continuous measurement of fluorescence. Data was collected using the CFX Maestro Software (BioRad, Hercules, California, USA) and analyzed using the 2^−ΔΔCT^ method [45].

### SH-SY5Y cells proliferation assay

Immediately before changing the cell transfection medium, 100 ng/mL of EGF were added to the SH-SY5Y culture medium in the defined conditions for 3-5 min [46]. Transfection medium was changed to a cell medium without FBS. Immediately after changing the medium, 10 μM of 5-ethynyl-2’-deoxyuridine (EdU) was added to the cell medium to perform the proliferation assay. Cells were maintained for further 12h, in a total of 18h of transfection.

Cell proliferation was determined using the Click-iT^®^ EdU Alexa Fluor^®^ imaging kit (Invitrogen) according to the manufacturer’s protocol. Briefly, coverslips containing the cells monolayer were fixed with 4% paraformaldehyde (PFA) in PBS for 15 min. After washing twice with 3% BSA in PBS the cells were permeabilized with 0.5% Triton X-100 in PBS for 20 min, and again twice washed. EdU detection is performed by incubating the coverslips with a Click-iT^®^ reaction cocktail for 30 min while protected from light, followed by a wash with 3% BSA in PBS. For subsequent DNA staining, coverslips were incubated with 5 μg/mL Hoechst 33342 in PBS for 30 min. Finally, coverslips were directly mounted on microscope slides with Vectashield^®^ mounting media (Vector Labs). EdU and Hoechst double labelled SH-SY5Y cells were imaged in an Olympus IX81 motorized inverted microscope equipped with a LCPlanFl 20×/0.40 objective lens (Olympus, Iberia S.A.U., Lisboa, Portugal). For each condition and each biological triplicate, random images of 20 fields were taken and EdU-positive and Hoechst-positive nuclei were manually scored using the Fiji software [47]. The total number of positive cells in the well was estimated based on the area covered by the objective, and the percentage of cell proliferation was determined by calculating the number of EdU-positive cells against the number of Hoechst-positive cells.

### Immunocytochemistry assays

HeLa and SH-SY5Y cells grown on coverslips were fixed with a 4% PFA solution for 20 min and, upon three washes with PBS and permeabilized with a 0.2% gr X-100 in PBS solution [43]. Following this, cells were blocked for 30 min-1h with 5% BSA in PBS, then incubated with specific primary antibodies, and finally incubated with fluorophore conjugated-secondary antibodies. Preparations were washed with PBS, mounted with Vectashield^®^ mounting media with or without DAPI. Cells were visualized by epifluorescence microscopy under an Olympus IX-81 motorized inverted microscope (as above) [48], or by confocal microscopy using a LSM 510 META confocal microscope (Carl Zeiss Microimaging GmbH, Jena, Germany) and a 63× oil objective [49].

### Morphometric analyses of neuronal-like SH-SY5Y cells

For cell morphology/morphometric assays, EGF treatment included a brief pulse of 3-5 minutes before changing the transfection medium. When indicated, the EGFR PD 168393 inhibitor was added 1h before the EGF stimulation. After 24h of cell transfection, cells were washed twice with PBS, fixed and processed for immunocytochemistry. Morphometric analysis of cells was performed on ~30 random digitized images per sample, with an average of 140 transfected cells per biological replica of each condition being analyzed. Transfected cells were identified by GFP expression, measurements were performed on matching acetylated-tubulin fluorescent microphotographs. All the measurements were performed as previously described [43], using Fiji. Process scoring was divided as follows: 1) “Total”, all processes arising from a cell; 2) “>20 μm”, all processes longer than 20 μm; and 3) “>35 μm”, processes longer than 35 μm (around two times the cell body length). Data is expressed as the number of processes per cell.

### Texas Red-conjugated Transferrin staining and co-localization analysis

Endocytosis of Texas Red-conjugated Transferrin (Molecular Probes) was monitored in SH-SY5Y 24h transfected with the GFP vector (‘V’), and APP-GFP [49]. In order to eliminate endogenous transferrin, cells were further incubated for 30 min at 37°C with DMEM medium supplemented with 20 mM HEPES. Medium was exchanged for 500 μL fresh medium of equal composition but containing 1mg/mL BSA and 100 nM Texas Red-conjugated transferrin, and cells incubated for a further 15 min at 37°C. The plates were immediately cooled to 4°C, washed twice with ice-cold PBS, and cells fixed and processed for immunocytochemistry. Cells were visualized by confocal microscopy in a LSM 510 META confocal microscope (Zeiss) and a 63× oil objective. To determine the percentage of co-localization between pro-EGF/EGF and Texas Red-transferrin, stack images of 20 delimited single cells (APP-GFP or control GFP vector transfected cells; from two independent experiments) were analyzed using the Fiji plugin JaCoP [50], and the percentage of co-localization of one protein with another was obtained by applying the Manders’ method [51].

### Statistical analysis

Data are expressed as mean ± SEM (standard error of the mean). The number of independent experiments (‘n’) and the test applied to infer statistical significance are stated in each figure caption. Data normality was tested by the Shapiro-Wilk (Royston method). Statistical analysis of normal distributed data was performed using the unpaired two-tailed Student’s t-test (to compare 2 groups), or the one-way ANOVA (to compare >2 groups) followed by the Tukey-Kramer multicomparison post-tests. Non-normally distributed data were analyzed by the Mann Whitney test (to compare 2 groups), or by the Kruskal-Wallis followed by the Dunn’s multicomparison post-test (to compare >2 groups). All statistical analyses were performed using the GraphPad Prism 8 software.

## Results

### APP binds to the HB-EGF protein

The HB-EGF (insert size 2.2-2.3 kb) was obtained as a positive clone from two distinct large-scale APP yeast two-hybrid screens performed by our group [52]. This interaction was validated by yeast co-transformation using two Gal4 DNA-BD bait plasmids, the pAS2-1-APP and the pAS2-1-AICD. This latter was used to test whether the interaction was mediated by AICD, the APP intracellular domain. Autoactivation tests confirmed that both the bait plasmids and the prey plasmid (Gal4-AD pACT2-HB-EGF) are unable to activate the reporter genes HIS3, ADE2 and MEL1, and to grow when independently transformed into the AH109 yeast strain cultivated in selective His^−^/Ade^−^ media. In the yeast co-transformation assay, prey and bait plasmids co-transformants were tested in SD/TDO, SD/QDO and SD/QDO/X-α-Gal medium. As shown in Fig 1a, pAS2-1-APP/pACT2-HB-EGF co-transformed bacteria were able to grow on SD/QDO (left image) and on SD/QDO/X-α-Gal media (right image, blue colonies on the top) with an appearance similar to the positive control (‘pvA3-1+pTD1-1’, right image). On the other hand, no growth of pAS2-1-AICD/pACT2-HB-EGF co-transformed colonies were verified, as in the negative control (pAS2-1+pACT2, right image). These yeast co-transformation assays confirmed that HB-EGF interacts with full-length APP, and that this interaction is independent of the APP AICD domain.

**Figure 1.**
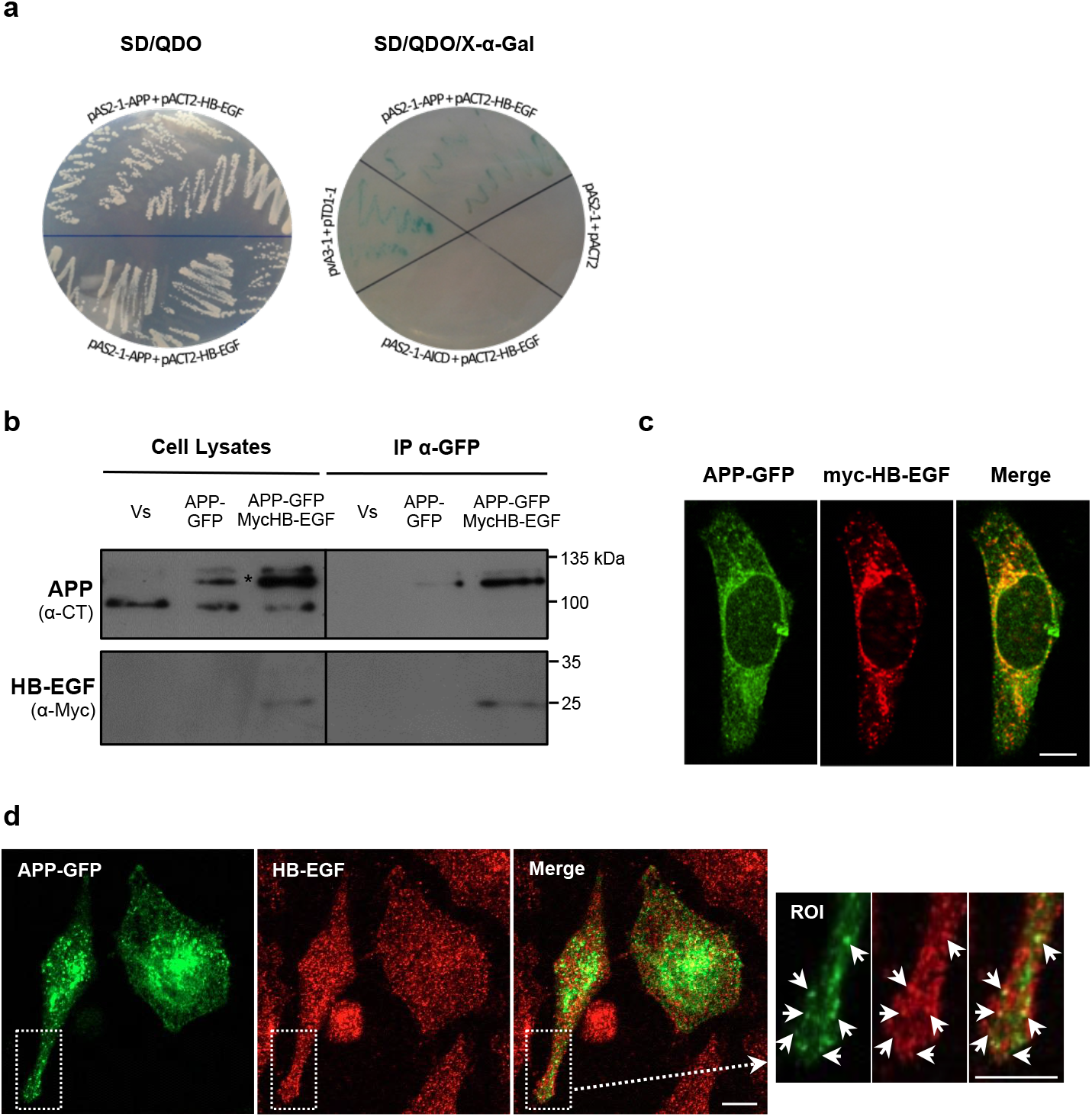
APP interaction with HB-EGF in yeast and mammalian cells. **a** Confirmation of the interaction between APP and HB-EGF by yeast co-transformation. Growth of yeast cells co-transformed with APP and HB-EGF constructs (pACT2-HB-EGF+pAS2-1-APP) was analyzed by streaking into quadruple dropout medium (QDO; -Leu, -Trp, -His, -Ade) in the absence of X-α-Gal (left). Yeast cells co-transformed with pACT2-HB-EGF+pAS2-1-APP, pACT2-HB-EGF+pAS2-1-AICD, and with the positive (pvA3-1+pTD1-1) and negative (pAS2-1+pACT2) controls were grown on QDO medium with X-α-Gal to test for α-galactosidase activity (blue colonies; right side). **b** APP interaction with proHB-EGF in HeLa cells. APP/HB-EGF interaction was validated in human HeLa cells transiently co-overexpressing the GFP and Myc empty vectors (‘Vs’), the Myc vector and the APP695-GFP cDNA (‘APP-GFP’), or the APP695-GFP and the Myc-HB-EGF cDNAs, which were immunoprecipitated using GFP-Trap^®^ magnetic beads. APP and the Myc-HB-EGF proteins were detected in cell lysates and in immunoprecipitates of APP-GFP transfected cells with anti-APP C-terminus and anti-Myc antibodies. Asterisk-marked bands represent mature and immature exogenous APP-GFP forms. **c** APP-GFP (green) co-localizes with exogenous Myc-HB-EGF (labelled with anti-Myc antibody, in red) at the perinuclear region (potentially the Golgi) and cytoplasmic vesicles (yellow/orange fluorescence in Merge). **d** Endogenous HB-EGF proteins (immunolabeled with an anti-HB-EGF antibody, red) co-localize with transfected APP-GFP (green) particularly at small vesicles distributed throughout the cytoplasm. ROI, region of interest. Bar = 5 μm.

To validate the APP-HB-EGF interaction in human cells, the GFP-trap technique was performed. HeLa cells were transiently transfected with APP695 tagged with GFP (‘APP-GFP’) alone or together with Myc-HB-EGF. Cells co-transfected with both GFP and Myc empty vectors were used as control (‘Vs’). APP-GFP was immunoprecipitated together with its interactors (Fig. 1b, ‘APP’ blot). Immunoblot using the anti-Myc antibody showed that the exogenous Myc-HB-EGF, which appeared around 25 kDa as expected, coimmunoprecipitated with APP-GFP in co-transfected cells (Fig. 1b, ‘APP-GFP MycHB-EGF’ lane in IP α-GFP). As expected, no APP was pulled down in the control condition (Fig. 1b, ‘Vs’ in IP α-GFP).

Immunocytochemistry studies were performed to address potential subcellular regions of APP/HB-EGF interaction. First, the subcellular distribution of the Myc-HB-EGF construct was analyzed and compared to the endogenous HB-EGF one (Online Resource 1). HB-EGF has been described to traffic between the plasma membrane (PM) and endoplasmic reticulum (ER), and can be found at the ER, Golgi apparatus, PM, nuclear envelope and perinuclear vesicles [53]. Accordingly, endogenous HB-EGF (red fluorescing, labeled with the anti-HB-EGF antibody) was distributed throughout the entire cell in vesicles and membrane compartments, slightly enriched at the perinuclear region. The Myc-HB-EGF fusion protein (green fluorescing, detected by the anti-Myc-tag antibody) localizes predominantly at the perinuclear region and nearby cytoplasmic vesicles, where it co-localizes with the endogenous HB-EGF (Online Resource 1).

The subcellular distribution of APP-GFP and both exogenous Myc-HB-EGF and endogenous HB-EGF, was compared in APP-GFP transfected cells. Micrographs show the expected distribution of transfected APP-GFP [40, 48, 49], enriched in organelles such as the ER, Golgi apparatus and cytoplasmic vesicles, and with a small fraction being present at the PM. APP-GFP and Myc-HB-EGF showed a considerable degree of co-localization at the perinuclear region, presumably the Golgi apparatus and nearby vesicles (Fig. 1c). Further, APP-GFP also co-localized with endogenous HB-EGF, predominantly at small vesicles distributed throughout the cytoplasm (Fig. 1d).

### APP interacts with pro-EGF

HeLa cells co-overexpressing APP-GFP and Myc-HB-EGF were used to pull down the APP-GFP protein and its interactors via the GFP-trap technique (Fig. 2a). The high amounts of immunoprecipitated APP-GFP (asterisk in Fig. 2a) also pull-downed endogenous APP (arrow in Fig. 2a), confirming that these proteins dimerize [54, 55]. Immunoblot with the anti-Myc antibody showed that the Myc-HB-EGF again was highly immunoprecipitated with APP-GFP (Fig. 2a, ‘APP-GFP/Myc-HB-EGF’ lane in IP α-GFP). Immunoblot analysis using the anti-HB-EGF antibody detected the recombinant Myc-HB-EGF and three faint protein bands between 33 and 37 kDa, found in all cell lysates. These immunoreactive bands most probably correspond to unmodified and post-translationally modified pro-HB-EGF forms [18, 56–58]. Although very faintly, all these bands were apparently detected in the APP-GFP pull-downs, suggesting that APP also interacts with endogenous pro-HB-EGF (Fig. 2a, IP α-GFP).

**Figure 2.**
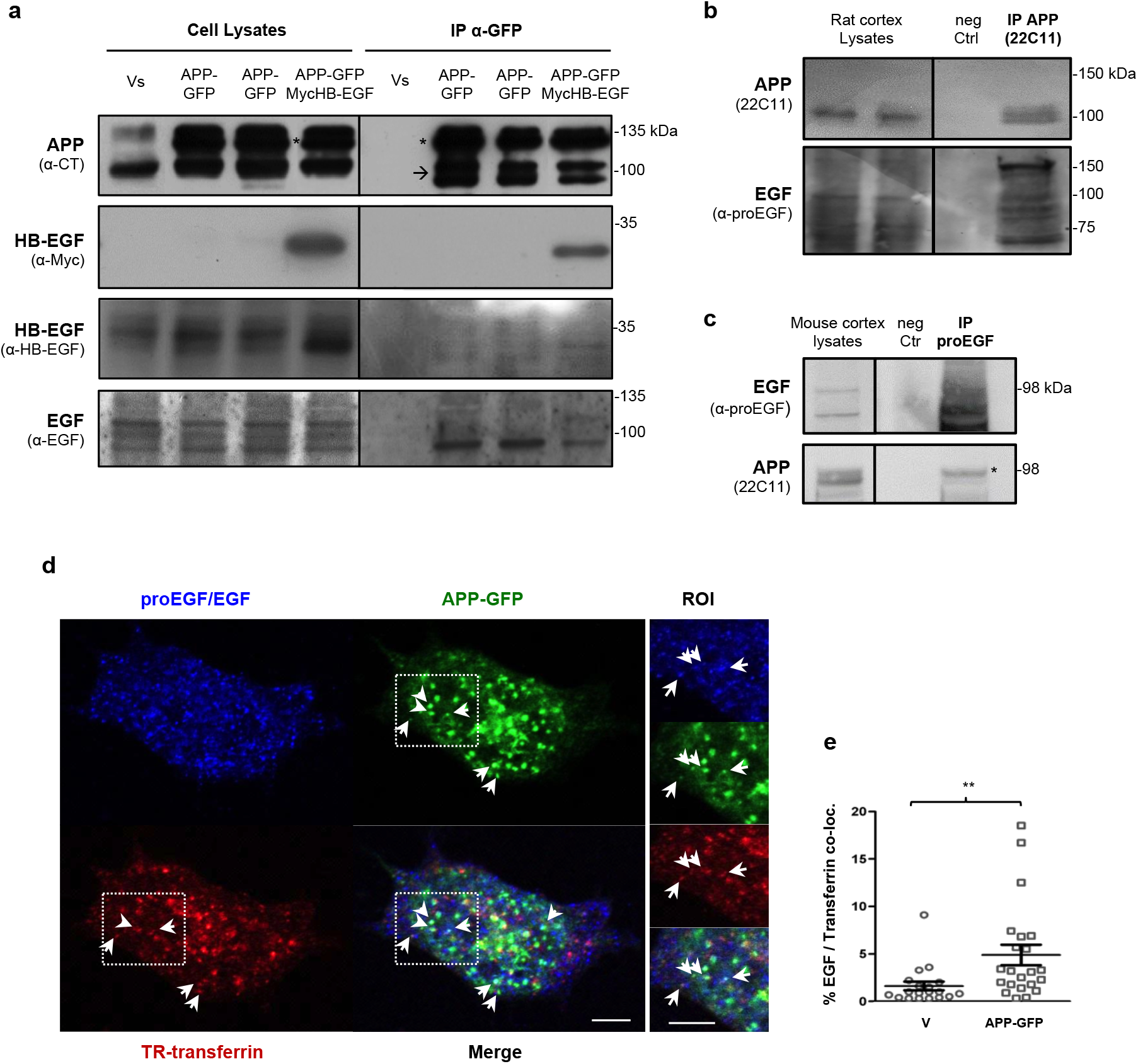
APP interacts with precursor forms of EGF and HB-EGF. **a** Lysates of HeLa cells transiently co-overexpressing the GFP and Myc empty vectors (‘Vs’), the Myc empty vector and the APP695-GFP cDNA (‘APP-GFP’), or the APP695-GFP and the HB-EGF cDNAs, were immunoprecipitated using GFP-Trap^®^ magnetic beads. Endogenous APP (arrow in blot) and transfected APP-GFP (asterisk in blot) were detected with an anti-APP C-terminal antibody; the anti-Myc-tag antibody detected the Myc-HB-EGF protein (~33 kDa); the anti-HB-EGF antibody detected various forms of endogenous HB-EGF (pre-pro-HB-EGF, at 37, 33, and 31 kDa); the anti-EGF antibody revealed different precursor EGF forms (94-130 kDa), including one of ~94 kDa that highly co-immunoprecipitated with APP-GFP. **b** Rat cortices were lysed with CHAPS buffer and immunoprecipitated with the anti-APP N-terminus 22C11 antibody. Immunoblotting with anti-proEGF antibody (specific for the EGF precursor form) revealed several immunoreactive pro-EGF bands in cortex lysates, that co-precipitated with APP. **c** Mouse cortices were lysed with NP-40 buffer and immunoprecipitated with anti-pro-EGF antibody. Immunoblotting with anti-APP (22C11) showed that APP co-precipitated with pro-EGF. Negative control (‘neg Ctr’): extracts incubated with protein G Dynabeads only (no primary antibody). **d** Confocal micrographs (plasma membrane focus plane) showing some co-localization between cellular pro-EGF/EGF (blue), endocytosed transferrin (red) and APP-GFP (green). SH-SY5Y cells transfected for 24h with APP-GFP cDNA were incubated at 37°C for 15 min with Texas-Red conjugated transferrin to allow for transferrin endocytosis. This is a time point upon which endocytosed transferrin is expected to be at the early endocytic route. Upon fixation, cells were immunolabeled for endogenous pro-EGF/EGF. Arrows indicate cytoplasmic vesicles at which pro-EGF, APP-GFP and transferrin appear to co-localize. ROI, region of interest. Bar = 5 μm. **e** EGF/Texas-red Transferrin co-localization (by Manders’ method using the Fiji plugin JaCoP) increases in cells expressing APP-GFP for 24h, in comparison with cells transfected with the GFP empty vector (‘V’). n = 20 cells per condition, from two independent experiments. **p < 0.01 by the Mann Whitney test.

Since EGF belongs to the same family of growth factors as HB-EGF, and due to the display of overlapping EGF cell and tissue expression and functions with APP, a physical interaction between APP and EGF forms was tested. The anti-EGF antibody detected 3 protein forms of pro-EGF in HeLa cells lysates (~95 to 120 kDa, possibly different glycosylated forms), with the lowest molecular weight band (~95 kDa) co-immunoprecipitating with APP-GFP (Fig. 2a, all APP-GFP lanes in IP α-GFP). The pro-EGF band immunoprecipitating with APP is fainter in APP-GFP and Myc-HB-EGF co-transfected cells, suggesting an EGF/HB-EGF competition for APP binding. The physical interaction between APP and pro-EGF was further analyzed in rat and mouse cortex lysates. Rat cortex lysates, homogenized and lysed with a non-denaturant CHAPS buffer, were immunoprecipitated with anti-APP N-terminal antibody 22C11. Figure 2b shows that the majority of the pro-EGF immunoreactive bands found in rat cortex lysates are co-precipitated with APP. In mouse cortex, lysed with a non-denaturant NP-40 buffer and immunoprecipitated with the anti-pro-EGF antibody, pro-EGF appeared as 3 immunoreactive bands: one around 98 kDa and other two close bands below this marker (Fig. 2c, ‘EGF’ blot). Concordantly with the above results from HeLa cells and rat cortex, pro-EGF was able to co-immunoprecipitate full-length APP. The APP form of higher molecular weight, most probably corresponding to its mature glycosylated form, is particularly more efficiently co-precipitated with pro-EGF (Fig. 2c, ‘*’ in ‘APP’ in IP proEGF). Pro-EGF could also be co-precipitated with an anti-APP antibody in mouse cortex lysates (data not shown). Of note and as expected, no APP was pulled down in any of the immunoprecipitation control conditions (‘Vs’ in Fig. 2a and ‘neg Ctrl’ in Fig. 2b and c).

Subsequent analysis of the subcellular co-distribution of endogenous pro-EGF/EGF and transfected APP-GFP revealed some degree of co-localization at cytoplasmic vesicles near the PM (Fig. 2d). Some of these proved to be endocytic, co-localizing with endocytosed fluorescent transferrin (arrows in Fig. 2d ROI). The percentage of co-localization between pro-EGF/EGF and endocytosed fluorescent transferrin was scored and observed to increase in conditions of APP-GFP overexpression, when compared to GFP expressing cells (Fig. 2e).

### Pro-HB-EGF and EGF increase APP protein levels

To study the effect of pro-HB-EGF on APP protein levels, HeLa cells were transfected with Myc-HB-EGF and the APP-GFP cDNAs, alone or together, or with the corresponding empty vector (‘Vs’). Cells expressing Myc-HB-EGF almost doubled the endogenous APP protein levels of V samples (Fig 3a and b, ‘endogenous APP’). This increase occurs for both immature and mature APP forms. We could not observe this increase in exogenous APP-GFP (Fig 3a and b, ‘APP-GFP’), potentially due to an artifact of the co-transfection itself (lowers amount of each co-expressed exogenous fusion protein). Another explanation could reside in the fact that HB-EGF increases transcription and translation of the *APP* gene itself. To pursue this, HeLa cells were transfected with the Myc-HB-EGF construct, and the levels of endogenous APP mRNA monitored by real time PCR. Although with some variability, Myc-HB-EGF overexpression did not alter *APP* gene expression (Fig. 3c). Of note, no alterations were observed for secreted sAPP levels in the Myc-HB-EGF transfected cells (Fig. 3a, ‘sAPP’).

**Figure 3.**
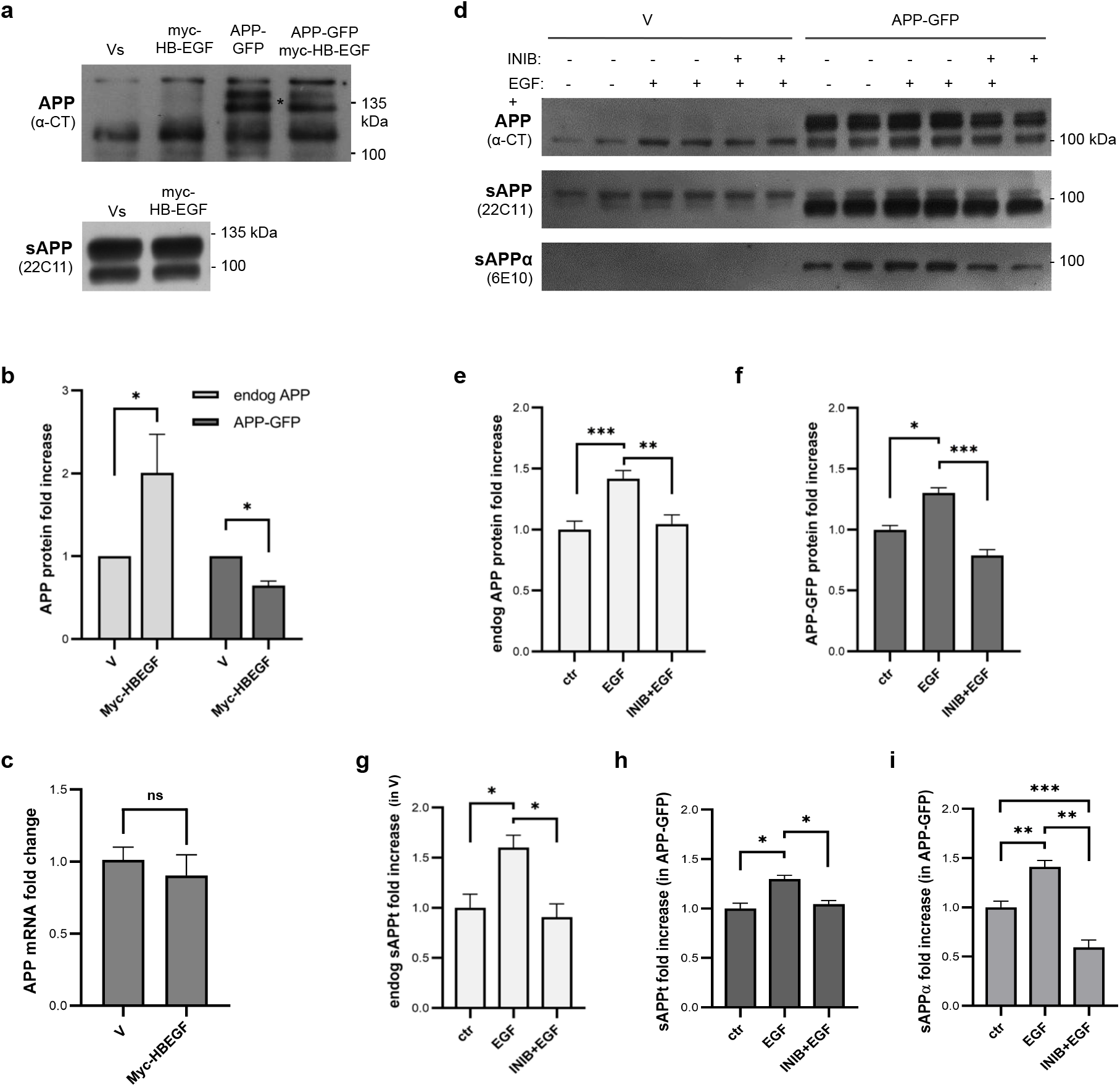
HB-EGF and EGF effects on APP protein levels and processing to sAPP. **a** Myc-HB-EGF expression in APP and sAPP protein levels. HeLa cells were co-transfected for 24h with the following cDNAs: the GFP and Myc empty vectors (‘Vs’), the GFP empty vector and the Myc-HB-EGF cDNA (‘HB-EGF’), the Myc empty vector and the APP695-GFP cDNA (‘APP-GFP’), or the APP695-GFP and the HB-EGF cDNAs. Immunoblots of cell lysates and medium were probed with anti-APP C-terminal (‘APP CT’) and N-terminal (‘22C11’) antibodies. Asterisk-marked bands represent exogenous APP-GFP bands. **b** Immunoblot data was plotted as fold increases of endogenous and exogenous (APP-GFP) APP in Myc-HB-EGF transfected cells against control conditions (established as 1.0). n = 3. *p<0.05, **p<0.01 and ***p<0.001 using the Student’s t-test. **c** Levels of endogenous APP mRNA in Myc empty vector (‘V’) and Myc-HB-EGF transfected cellular lysates were monitored by real time PCR. Data is plotted as a fold increase of control empty vector. n = 4. Statistically non-significant results by the Student’s t-test. **d** The effects of EGF incubation in APP and sAPP levels were assessed in SH-SY5Y cells, either transfected with the GFP empty vector (‘V’) or the APP695-GFP (‘APP-GFP’) cDNAs, and stimulated or not for 3 min with 1OO ng/mL human recombinant EGF (‘V+EGF’ and ‘APP+EGF’) in the presence (‘V+IN+EGF’ and ‘APP+IN+EGF’) or not of 1O μM of the EGFR inhibitor PD168393. Cellular lysates were probed with the anti-APP C-terminal (‘APP CT’), and media lysates with the anti-APP N-terminal 22C11 (for total soluble APP, ‘sAPP’) and the anti-APP 6E1O (for ‘sAPPα’) antibodies. **e** Endogenous APP (‘endog APP’) full-length protein levels were plotted as fold increases of the respective non-treated controls. n = 12. **p < 0.01, ***p < 0.001 by one-way ANOVA followed by the Tukey’s multicomparison post-test. **f** Exogenous APP-GFP full-length protein levels were plotted as fold increases of the non-treated APP-GFP overexpressing cells. n = 9. *p < 0.05, ***p < 0.001 by Kruskal-Wallis followed by the Dunn’s multicomparison post-test. Of note, ctr vs INIB+EGF has p=O.OO56 by the Mann Whitney test. **g** Total sAPP protein levels were monitored in the cell media from non-transfected cells (‘endog sAPPt’). **h** Total sAPP protein levels were monitored in the cell media from APP-GFP transfected cells (‘sAPPt in APP-GFP’). **i** sAPP resulting from the alpha-secretase cleavage (‘sAPP alpha’) was also determined in APP-GFP transfected cells. All quantifications of sAPP levels (g-i) were plotted as fold increases of the respective non-treated cells. n = 6. *p < 0.05, **p < 0.01, ***p < 0.001 by one-way ANOVA followed by the Tukey’s multicomparison post-test. Ponceau S staining of total proteins bands was used as loading control for all the western blot data, and data is presented as mean ± standard error of the mean.

Since EGF stimulation has been reported to regulate APP processing, we further confirmed this to occur under our experimental conditions (Fig. 3d-i). SH-SY5Y cells overexpressing the GFP empty vector (‘V’) or APP-GFP for 24h, were exposed (‘EGF +’) or not (‘EGF -’) to a 3 min EGF pulse in the presence (‘INIB +’) or absence (‘INIB -’) of the EGFR inhibitor, incubated for 1h, 17h before medium and cell harvesting. Immunoblots of cell lysates show that EGF stimulation significantly increased both endogenous APP and exogenous APP-GFP levels. Concomitant EGFR inhibition blocked this effect (Fig. 3d-f), and even decreased exogenous APP-GFP levels below control ones (Fig.3d and f; p=0.0056 by the Mann Whitney test). Furthermore, EGF also significantly stimulated total sAPP (‘sAPPt’) secretion, either in control cells (‘endog sAPPt’; mainly sAPP_751/770_) or in APP-GFP expressing cells (‘sAPPt in APP-GFP’, both endogenous sAPP_751/770_ and exogenous sAPP_695_) (Fig. 3d ‘sAPP’, and 3g-h). In APP overexpressing cells, EGF also stimulated sAPPα release, detected by the 6E10 antibody (Fig. 3d ‘sAPPα’, and 3i ‘sAPPα in APP-GFP’). As seen for cellular APP protein levels, EGFR inhibition blocked the EGF-induced sAPP production/secretion. Noteworthy, EGFR treatment in APP overexpressing cells significantly decreased sAPPα levels below control cells levels.

### APP effects on basal and EGF-induced ERK activation

Figure 3 results showed that EGF stimulation increases APP protein levels and sAPP release in SH-SY5Y cells, in an EGFR activation dependent manner. EGFR activation by EGF can trigger downstream signaling pathways, particularly ERK1/2 activation. APP effects on basal and EGF-induced ERK1/2 activation were thus monitored. The levels of total and phosphorylated ERK1/2 (‘pERK’) were analyzed in both HeLa and SH-SY5Y cells overexpressing for 24h the GFP empty vector (‘V’) or the APP-GFP (‘APP’), exposed or not for 30 min to EGF before cell harvesting (Fig. 4, ‘pERK’ and ‘tERK’). In both cell lines, and as expected, EGF treatment alone greatly increased the phosphorylated ERK1/2 levels and the pERK/ERK ratio (Fig. 4, ‘V+EGF’ vs ‘V’). APP-GFP overexpression alone induced a modest ~2-fold increase in the pERK1/2/ERK ratio (Fig. 4f, p=0.0043 by the Mann Whitney test for ‘APP’ vs ‘V’ in SH-SY5Y cells). In this experimental design, of 24h of APP-GFP expression and EGF stimulation in the last 30 min, combination of APP-GFP overexpression and EGF treatment synergistically promoted ERK1/2 activation, significantly in HeLa cells (Fig. 4a-c, ‘APP+EGF’ vs ‘V+EGF’).

**Figure 4.**
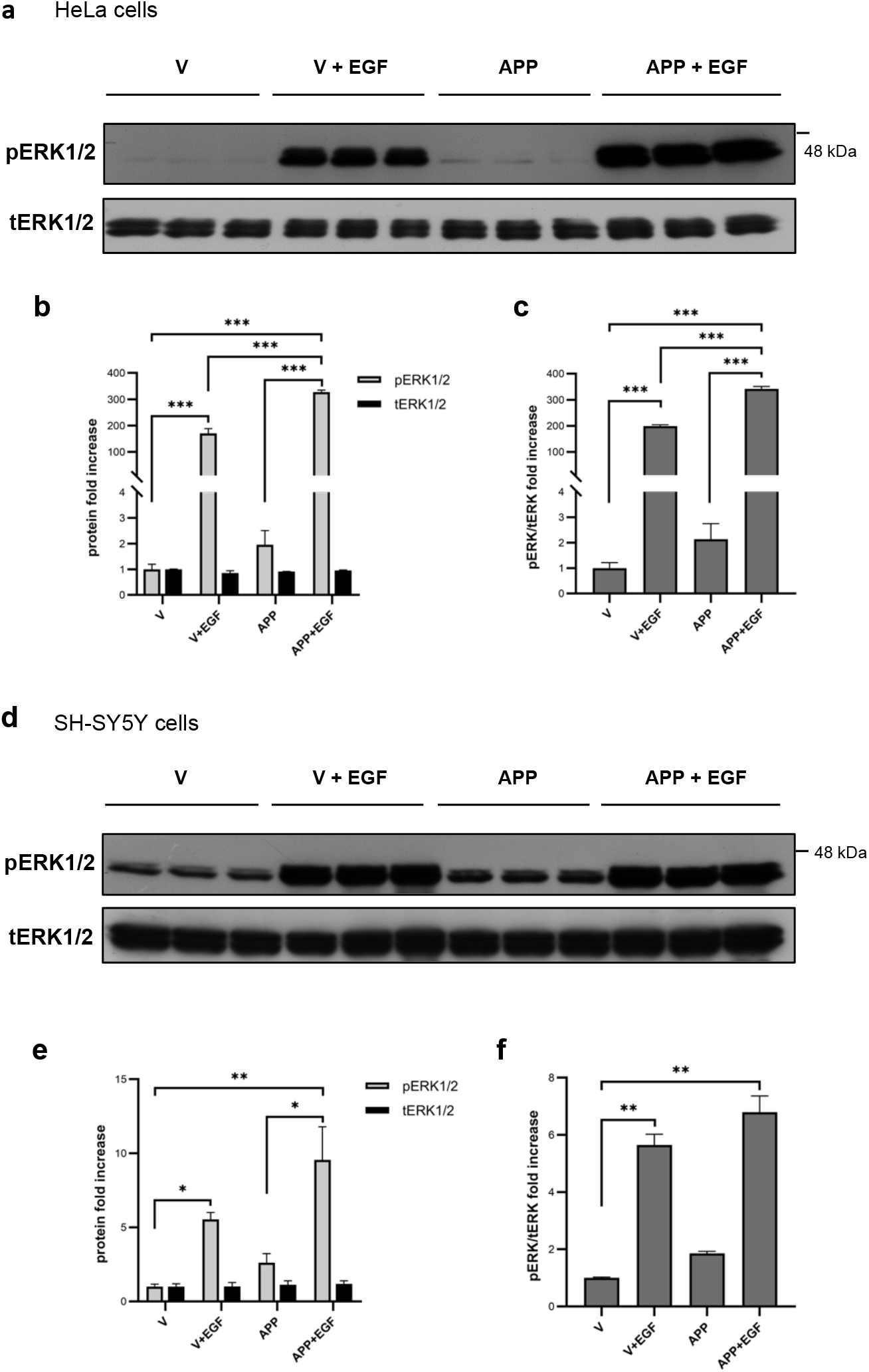
APP effects on basal and EGF-induced ERK signaling in HeLa and SH-SY5Y cells. Cells were either transfected with the GFP empty vector (‘V’) or the APP695-GFP cDNA (‘APP’) for 24h, and treated or not with 100 ng/mL human recombinant EGF (‘V+EGF’ and ‘APP+EGF’) 30 min before cell harvesting. **a** Immunoblots of HeLa cell lysates probed with the phospho-ERK1/2 (‘pERK1/2’), and total ERK 1/2 (‘tERK1/2’) antibodies. **b** HeLa cells’ pERK1/2 and tERK1/2 protein levels. **c** HeLa cells’ pERK/tERK ratio. n = 3. ***p < 0.001 by one-way ANOVA followed by the Tukey’s multicomparison post-test. Protein levels were plotted as fold increases of control empty vector (“V”). **d** Immunoblots of SH-SY5Y cell lysates probed with the phospho-ERK1/2 (‘pERK1/2’), and total ERK 1/2 (‘tERK1/2’) antibodies. **e** SH-SY5Y cells’ pERK1/2 and tERK1/2 protein levels. **f** SH-SY5Y cells’ pERK/tERK ratio. n = 5-6. * p < 0.05, **p < 0.01 by Kruskal-Wallis followed by the Dunn’s multicomparison post-test. Protein levels were plotted as fold increases of control empty vector (“V”). Ponceau S staining of total proteins bands was used as loading control for all the data, and data is presented as mean ± standard error of the mean.

### APP potentiates EGF-mediated neuritogenesis in SH-SY5Y cells

Following the evidence on the physical interaction between APP and pro-EGF, the role of such interaction was pursued. Since EGF is a known mitogen, we investigated if APP could alter EGF-mediated proliferation in neuroblastoma cells (Online Resource 2). To address this issue, we performed a proliferation assay based on the incorporation of EdU, a nucleoside analog of thymidine that is incorporated into DNA during cell division. As follows, SH-SY5Y cells were transfected with the GFP empty vector (‘V’) or with APP-GFP (‘APP’) for 18h, and exposed or not to a 3 min EGF pulse right before transfection medium exchange. After the EGF pulse, the DNA of SH-SY5Y cells undergoing mitosis was labelled with EdU (Online Resource 2a-b). In these conditions, no differences in the percentage of proliferating cells (EdU-positive/Hoechst-positive nuclei ratio) were observed between all the conditions (Online Resource 2c). Although EGF did not potentiate proliferation over basal values, at the assay’s endpoint there was a higher number of EGF-stimulated cells than non-stimulated cells, indicating an EGF pro-survival effect. APP had no influence in this EGF pro-survival effect, as no differences were observed between APP transfected cells and vector transfected cells (Online Resource 2d).

A closer look on the morphology of the SH-SY5Y cells at the end of the proliferation assay (experimental design in Fig. 5a), suggested that APP-EGF interaction could be key for neuritogenesis, since longer neuritic-like processes were observed in APP+EGF conditions, when compared to any other condition (Fig. 5b microphotographs). This putative functional interaction is corroborated by the parallel highest increase in ERK1/2 phosphorylation levels, a pathway known to be involved in neuronal differentiation [59–61] (Fig. 5, ‘APP+EGF’). To better elucidate this hypothesis, we performed several morphometric analyses in SH-SY5Y cells transfected for 24h with APP-GFP and stimulated at the 8^th^ hour of transfection with a 3-5 min pulse of EGF (Fig. 6a). Since EGFR activation is known to mediate EGF neuritogenic effects, we also studied the role of EGFR in the APP+EGF-induced neuritogenesis, by additionally pre-incubating cells with the EGFR inhibitor PD168393 (Fig. 6). Morphometric analyses included scoring the total number of neurite-like processes arising from each cell (‘Total’, all processes of all sizes), and measuring and categorizing these processes by length: processes higher than 20 μm (pre-neurites), and processes higher than 35 μm (neurites) [43].

**Figure 5.**
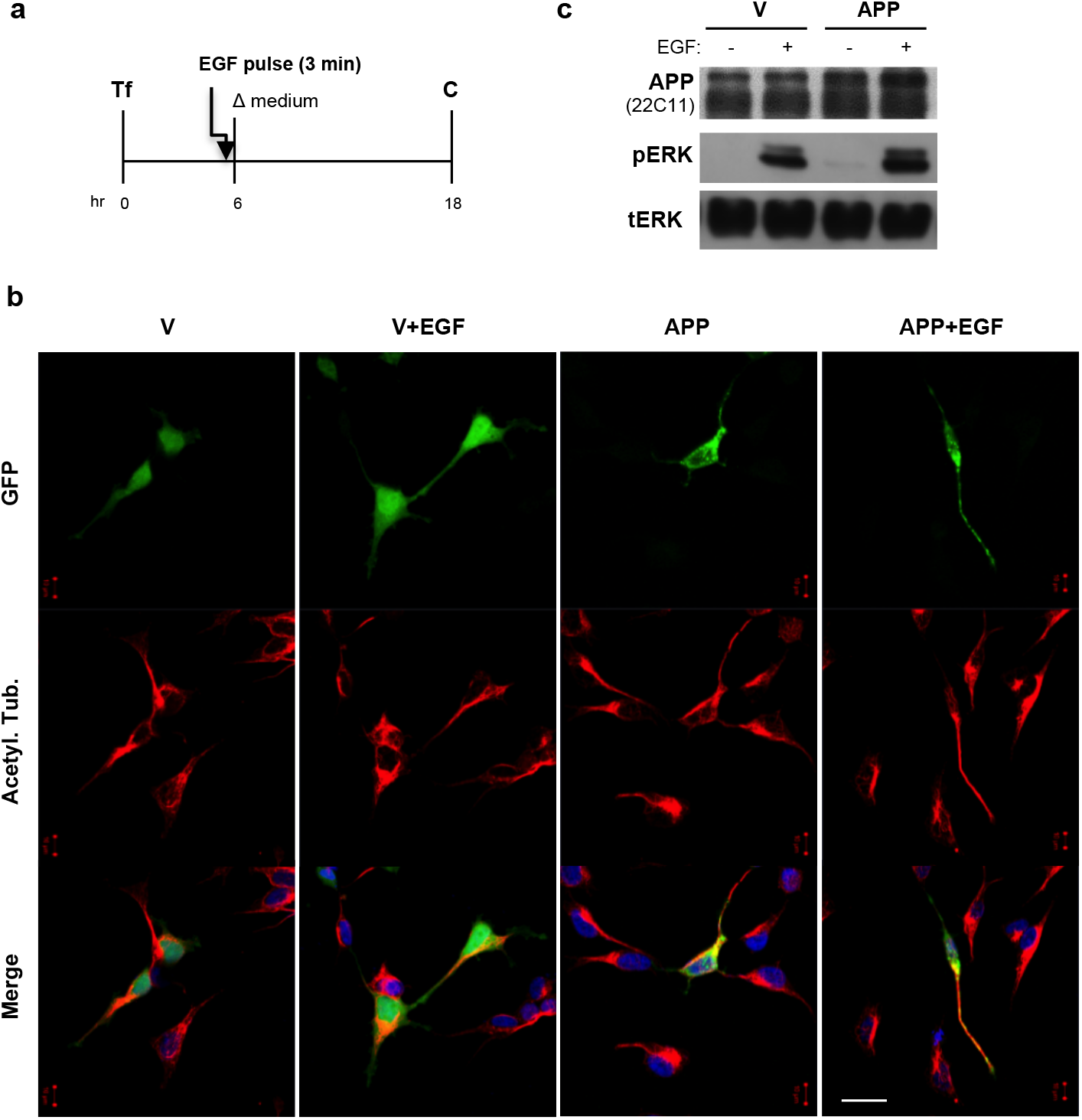
APP promotes EGF-induced neuritogenesis, potentially via ERK activation. **a** SH-SY5Y cells, either transfected (‘Tf’) with the GFP empty vector (‘V’) or APP695-GFP (‘APP’) cDNAs, were stimulated or not for 3 min with 100 ng/mL of human recombinant EGF (‘V+EGF’ and ‘APP+EGF’), 12hr before fixation/harvesting. **b** Representative epifluorescence microphotographs of these GFP or APP-GFP (green) expressing cells subjected to immunocytochemistry procedures to label the differentiation marker acetylated α-tubulin (‘Acetyl. Tub.’, in red), and nuclear stained with DAPI (blue). Bar = 20 μm. Cell morphology suggests a synergy between APP and EGF on SH-SY5Y neuronal-like differentiation. **c** Immunoblots of cell lysates, probed with the 22C11 antibody for the APP N-terminus, and for phospho-ERK1/2 (‘pERK’) and total ERK 1/2 (‘ERK’), suggest a synergy between APP and EGF on ERK1/2 activation.

**Figure 6.**
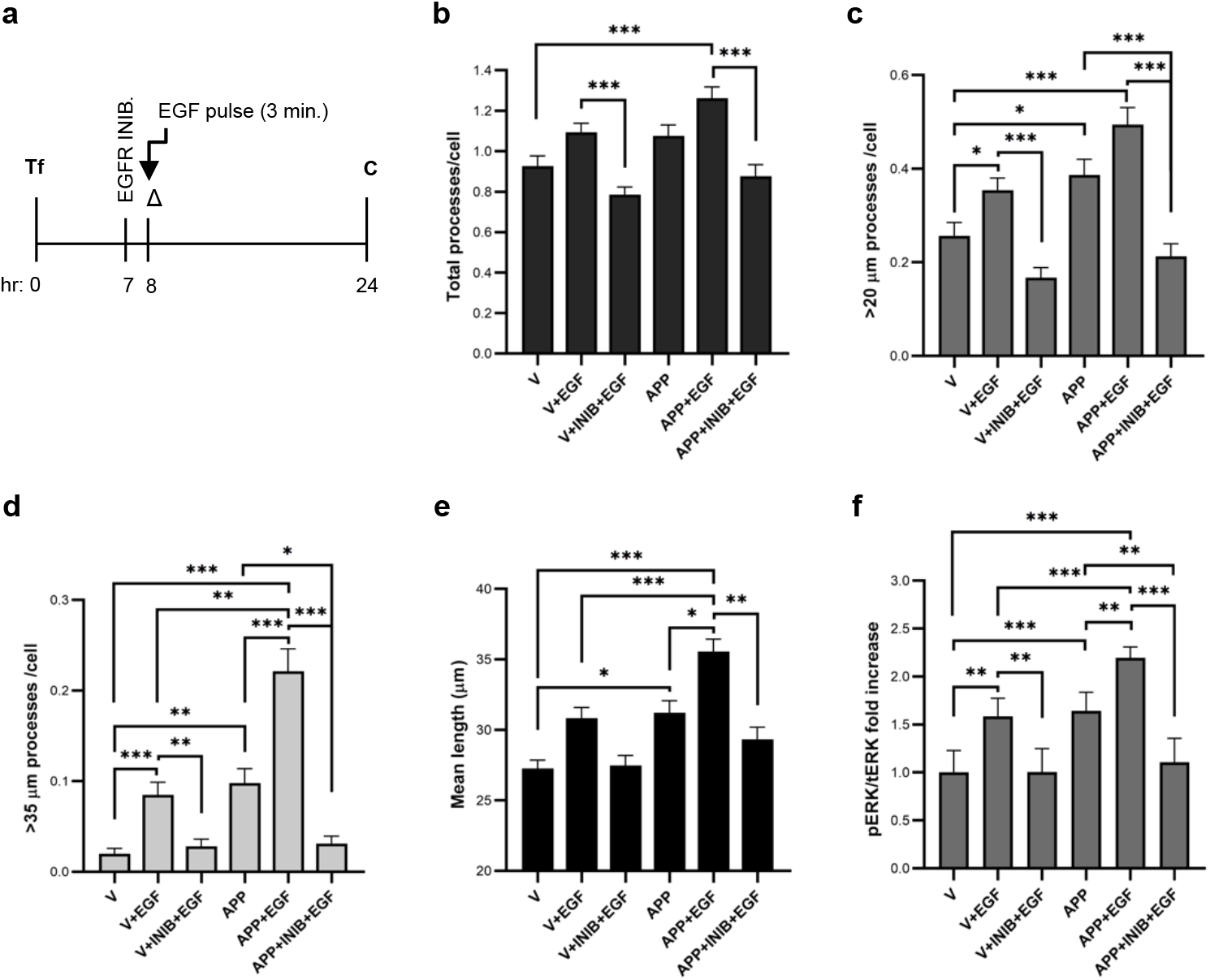
APP increases EGF-induced neuritic elongation in an EGFR/ERK-dependent manner. **a** SH-SY5Y cells, either transfected (‘Tf’) with the GFP empty vector (‘V’) or APP695-GFP (‘APP’) cDNAs, were stimulated or not for 3 min with 100 ng/mL of human recombinant EGF (‘V+EGF’ and ‘APP+EGF’) immediately before cell media change (16hr before fixation/harvesting). When indicated, this EGF stimulation was performed under EGFR inhibition with 10 μM PD168393 (‘INIB’), added 1h before EGF. Cells were fixed and collected for analysis after 24h of cell transfection. **b** Total number of processes per cell – all processes protruding from a cell, independently of the size, were scored. **c** Number of processes longer than 20 μm per cell. **d** Number of processes longer than 35 μm per cell. **e** Mean length of processes longer than 20 μm. n = 4; around 30 microphotographs, corresponding to ~100 cells per condition/assay, were analyzed. *p < 0.05, **p < 0.01, ***p < 0.001 by Kruskal-Wallis followed by the Dunn’s multicomparison post-test. Statistical differences, not fundamental for result interpretation, were omitted. **f** pERK/ERK ratio was calculated based on immunoblots of phospho-ERK1/2 and total ERK 1/2 and plotted as fold increases over control “V” levels. Ponceau S staining of total proteins bands was used as loading control. n = 5. *p < 0.05, **p < 0.01, and ***p < 0.001 by one-way ANOVA followed by the Tukey’s multicomparison post-test. All data is presented as mean ± standard error of the mean.

The total number of processes tended to increase in cells only exposed to EGF or transfected with APP (Fig. 6b). Co-treatment with both APP and EGF resulted in their significant increase from 0.93±0.05 (control ‘V’) to 1.26 ± 0.06 (‘APP+EGF’). Relative to the number of processes longer than 20 μm, EGF-stimulation and APP overexpression alone significantly increased this number (Fig. 6c), and the concomitant overexpression of APP with EGF stimulation (‘APP+EGF’) tended to even increase it. This is particularly true for the longer processes. Combined APP overexpression and EGF treatment significantly increased the number of processes longer than 35 μm per cell (0.22 ± 0.025) when compared to any of the other conditions (0.02 ± 0.01 for V, 0.08 ± 0.01 for EGF and 0.10 ± 0.02 for APP) (Fig. 6d). Further, the combined action of APP and EGF was the only condition able to significantly increase neuritic length (Fig. 6e), accompanied by an increase in neuronal βIII-tubulin protein levels (Online Resource 3a, ‘APP+EGF’).

Cells concomitantly treated with EGF and the EGFR inhibitor (‘INIB’) showed a significantly lower number of processes per cell than cells treated with EGF alone, back or even lower than control numbers (Fig. 6b-d). The number of total processes per cell only slightly decreased in APP+EGF cells treated with the EGFR inhibitor, and the number was very similar to the APP condition alone (Fig. 6b, ‘APP+INIB+EGF’ vs ‘APP’). However, EGFR inhibition had a more drastic effect on neurite elongation, significantly decreasing the number of > 20 μm and > 35 μm processes, when compared to both ‘APP’ and ‘APP+EGF’ conditions (Fig. 6c-d). Further confirmation was obtained from the significant decrease in the mean processes’ length when APP overexpressing cells were exposed to EGF in the presence of the EGFR inhibitor (Fig. 6e, ‘APP+INIB+EGF’ vs ‘APP+EGF’). This also resulted in a decrease in the neuronal βIII-tubulin protein levels compared to the EGF+APP condition (Online Resource 3a, ‘APP+EGF+INIB’).

In the previous Figures 4 and 5 we showed that APP overexpression was associated with higher levels of phosphorylated ERK1/2 in human cell lines, in both basal and EGF-stimulated conditions. To confirm that ERK was a mediator of this APP/EGF combined neuritogenic effect via EGFR, ERK1/2 activation was monitored via WB means (Fig. 6f and Online Resource 3b, pERK1/2 and tERK1/2). At this time point, EGF-stimulated cells and APP-GFP transfection similarly increased the levels of pERK, and EGF stimulation in combination with APP overexpression further promoted the ERK activation state. The presence of the EGFR inhibitor hindered this EGF- and APP-induced pERK1/2 activation, even below APP only transfected cells. Noteworthy, we also analyzed phosphorylated STAT3 levels and, although EGF stimulation induced pSTAT3 via EGFR activation, APP overexpression had no further effect on these (data not shown).

Taken together, all these results indicate that neuritogenesis mediated by APP+EGF (Fig. 5a and 6b-e) involves ERK1/2 activation (Fig. 5b and 6f), in an EGFR activation-dependent manner (Fig. 6b-f), and might involve EGF-induced APP cleavage into sAPPα (Fig. 3i), the most neuritogenic sAPP fragment [62].

## Discussion

APP is a central protein in AD pathophysiology and has important neurodevelopmental and neuromodulatory roles. In the present work we studied the interaction between APP and two members of the EGF family of growth factors, HB-EGF and EGF.

HB-EGF was here confirmed as a novel APP interactor, after being previously identified by us as a putative APP interactor in a YTH screen of a human brain cDNA library. The interaction was validated by yeast co-transformation assays and by an APP-GFP/MycHB-EGF co-immunoprecipitation assay in a human cell line (Fig. 1 and 2).

After synthesis of the HB-EGF precursor, or readily as it reaches the cell surface, the HB-EGF propeptide domain is rapidly cleaved. PM-anchored HB-EGF mainly consists of its mature, transmembrane and cytoplasmic domains (commonly referred as pro-HB-EGF) [17]. Since the Myc tag of our Myc-HB-EGF cDNA construct is N-terminally located, detection of an APP-interacting band immunoreactive with the anti-Myc antibody indicates that APP is able to interact with the pre-pro-HB-EGF. A positive although faint interaction was also apparent between APP and HB-EGF endogenous species ranging from 31 to 37 kDa and detected by an anti-HB-EGF antibody. These bands most likely represent unmodified and glycosylated pre- and pro-HB-EGF species (predicted to have 18-26 or 26-34 kDa) [18, 56–58].

The yeast co-transformation assay shows that HB-EGF does not interact with the AICD, suggesting that the interaction must occur via the APP N-terminal. Two APP N-terminal domains, described to mediate APP growth and cell adhesion promoting activities might be responsible for APP binding to pro-HB-EGF: the cysteine-rich GFLD at E1 and the RERMS-containing GFLD at E2 [1, 63]. In fact, APP extracellular domain mediates its interaction with various extracellular matrix (ECM) components (like heparan sulfate proteoglycans, collagen type I, collagen type IV and laminin), and other cell adhesion molecules (NCAM and NgCAM) [6, 64]. Heparin binding to the APP E1 or E2 region can induce protein dimerization [1], and binding of APP to heparan sulfate proteoglycans (HSPG) is important for substrate bound-APP to induce neurite outgrowth [65]. Similar to APP, HB-EGF binds heparin and HSPG. HB-EGF interaction with cell surface HSPG, via its heparin-binding domain, is described to modulate HB-EGF binding and activity [17]. Moreover, PM-anchored HB-EGF forms complexes with other transmembrane proteins, including α3β1 integrin (Itg) and tetraspanins (e.g. CD9 and CD81) [66]. The interaction with CD9 modulates PM-anchored HB-EGF juxtacrine actions, cleavage to the mature form, and internalization [17, 66]. Interestingly, APP also forms complexes with α3β1 Itg and CD81 to mediate cell/growth cone adhesion [10, 67], and with α3β1 Itg and Reelin to mediate hippocampal dendritic outgrowth [9]. Thus, the formation of cis- and trans-heterodimers between APP and pro-HB-EGF might be mediated by HSPG, integrins and tetraspanins, and are expected to have a role in (neural) cell adhesion, migration and neuritogenesis.

The present work also proves that APP can form complexes with pro-EGF, including in the brain cortex (Fig. 2a-c). Pull downs of APP-GFP or endogenous APP were able to coimmunoprecipitate different pro-EGF forms. These pro-EGF forms may derive from its glycosylation on at least 10 glycosylation sites, from alternative splicing, and from pro-EGF prodomain cleavage [68]. Interestingly, lower levels of endogenous pro-EGF were coimmunoprecipitated with APP-GFP in cells expressing the Myc-HB-EGF (Fig. 2a), indicating that pro-EGF and pro-HB-EGF might compete for APP binding. This, together with the high levels of EGF in HeLa cells [69, 70], potentially explain the low rate of APP binding to endogenous HB-EGF in these cells.

Several studies show that EGF has neurotrophic and neuromodulatory functions [16, 24], and demonstrate its potential for neuroprotective and neuroregenerative strategies [71–73]. Such functions are mostly attributed to the mature soluble EGF molecule. Although there is evidence that in some tissues EGF accumulates as pre-pro-EGF and can release forms of different sizes [16, 19], there are very few studies exploring pro-EGF protein-protein interactions and its underlying functions. Soluble pro-EGF is described to bind to secreted modular calcium binding protein (SMOC), an ECM component and an antagonist of bone morphogenetic protein via ERK activation. SMOC was described to bind to pro-EGF but not to mature EGF, and is suggested to retain pro-EGF at the cell membrane by mediating pro-EGF binding to HSPGs [74]. Similarly, pro-EGF/APP/HSPGs complexes are likely to exist. Supporting our findings of APP interaction with pro-HB-EGF and pro-EGF, some known APP interactors have EGF-like domains. Namely, LRP1 [75, 76], SorLA [49, 77, 78] and NOTCH [79, 80], all proteins with well described effects on APP metabolism and that have been extensively implicated in the AD pathology. EGF-like domains are generally present in the extracellular domain of membrane-bound proteins or in secreted protein, which are often ECM proteins. Amongst these, the ECM proteins agrin [81, 82] and fibulin [83] may play a significant role in regulating APP neuronal differentiation functions [83–87]. The APP/fibulin interaction depends on the fibulin EGF-like domain and on the APP E1 region, and treatment with secreted fibulin blocks sAPP-stimulated neural stem cells proliferation [83]. Interestingly, fibulin also interacts with the extracellular domain of pro-HB-EGF [88], and fibulin/pro-HB-EGF/APP may form a macromolecular functional tricomplex.

APP binding to netrin-1 is dependent of the APP residues 1-17 of the Abeta domain and involves the netrin-1 EGF-like domain [89]. Since the EGF ligand itself (as well as other proteins bearing EGF-like domains) binds Abeta 42 fibrils and fibrillar aggregates [82], we speculate that the Abeta domain is also a probable APP binding site for pro-EGF and pro-HB-EGF. Future molecular biology experiments designed to understand the APP domain involved in these interactions will be valuable, as well as assays to test whether APP binds to other EGF-family growth factors and proteins with EGF-like domains.

Several of the above described APP interactors were observed to alter APP levels and processing [75, 76, 78, 81]. HB-EGF overexpression was here observed to induce an increase in total endogenous APP, either immature or mature (Fig. 3a-b). However, HB-EGF did not alter APP mRNA levels (Fig. 3c). Since the amount of the mature APP form was especially increased, we can postulate that HB-EGF is retaining mature APP in secretory vesicles and/or at the PM and is increasing APP half-life [49]. Similarly, EGF stimulation significantly increased APP full-length protein levels and secreted sAPP (both total and sAPPα); concomitant EGFR inhibition blocked both effects even below control levels for APP-GFP conditions (Fig. 3d-i). The mechanism behind EGF-EGFR induced sAPP secretion might include the activation of PKC activity and/or ERK1/2 (Fig. 5b and 6f) [34, 48, 49, 90, 91]. Since EGF induced an increase of both endogenous and exogenous APP, EGF could be promoting APP half-life, and not *APP* gene expression (as hypothesized for HB-EGF). We show that APP co-localizes with cellular pro-EGF/EGF in vesicles, some of endocytic nature, and that APP overexpression increases pro-EGF/EGF colocalization with endocytosed transferrin, suggesting that APP can modulate pro-EGF and EGF-EGFR endocytosis (Fig. 2d-e). In fact, we have collected data showing that APP also modulates EGFR expression levels and trafficking (da Rocha JF *et al*., in preparation). Concordantly, other studies have shown that APP up-regulation can induce Rab-5 overactivation and accelerate endocytosis [92, 93], and that APP can be trafficked in the same population of endosomes and multivesicular bodies that carry EGF-EGFR [94].

Using the neuronal-like SH-SY5Y cell model, we tested the existence of a functional interaction between APP and the mature EGF. No function in cell proliferation was observed for either EGF or APP in the conditions here tested (Online resource 2), and the mitogenic effect of EGF may not occur in SH-SY5Y cells as shown by others [95]. Nevertheless, EGF had a protective effect in stimulated cells, although co-treatment with APP did not increase EGF promotion of cell survival. Alternatively, morphometric analyses of SH-SY5Y cells 24h after APP overexpression and EGF treatment showed that this combination accelerates neurite outgrowth (Fig. 6 and Online Resource 3). Combined treatment significantly increases the number of processes per cell (including of longer processes), as well as the processes’ mean length. Although APP alone already significantly increases the cellular number of processes longer than 20 and 35 μm, compared to control empty vector, combination of APP with EGF further increases the number of processes longer than 35 μm per cell above all other conditions. This neuronal-like differentiation induced by combined APP and EGF treatment was accompanied by a lasting increase in ERK1/2 activation, detected even 16h after EGF stimulation. Accordingly, shifts in ERK1/2 signaling from transient to sustained were implicated in EGF-mediated neuronal differentiation [96, 97].

EGFR activation can trigger neurite outgrowth of neuroblastoma cells potential via PI3K-Akt and MEK-ERK1/2 activation [59, 61]. As expected, EGF treatment in the presence of an EGFR inhibitor decreased all the parameters to similar or slightly below control empty vector. Moreover, in APP overexpressing cells the EGFR inhibitor significantly decreased the number of >20 μm and >35 μm processes when compared not only to the APP+EGF condition but also to the APP overexpression alone, suggesting that EGFR activation is implicated (at least partially) in the role played by APP in neurite outgrowth. Results on the pERK/ERK ratio further support this conclusion. While stabilization of membrane-bound APP is important for its cell-adhesion functions [10], APP cleavage into sAPP is necessary to modulate neurite outgrowth [43, 98], and can be partially regulated by EGFR and ERK1/2 activation [90]. In accordance, the morphologic and signaling alterations here detected after EGF stimulation (Fig. 5 and 6) were accompanied by alterations in APP protein levels and in its processing to the neuritogenic sAPPα (Fig. 3d-i). The fact that APP activates ERK1/2, and cooperates with EGF-EGFR signaling, is suggestive of a positive-feedback pathway culminating in sAPP release, ERK1/2 activation and neurite outgrowth.

A synergistic interaction between APP and HB-EGF in neuritic outgrowth is also most likely to occur and should be pursued in APP-transfected primary neurons exposed to mature HB-EGF. Indeed, mature HB-EGF was already reported to promote neurite outgrowth in PC12 [28] and different neuroblastoma cell lines, including SH-SY5Y [99]. Induction of both neuritic outgrowth and decreased cell proliferation required HSPG and EGFR, and the downstream activation of ERK1/2 and STAT3 [99].

## Conclusions

Our study confirms pro-HB-EGF and identifies pro-EGF as novel APP interactors, in yeast and mammalian cells, and pro-EGF also in rodents’ brain tissue. APP overexpression and concomitant HB-EGF overexpression or exposure to EGF, both increase APP levels (potentially via increased APP half-life). We further show that EGF and APP synergistically activate the ERK signaling pathway, crucial for neuronal differentiation, and promote neuritogenesis, in a mechanism involving EGFR activation. Overall, this work reveals novel APP protein interactors bearing EGF domains and provides mechanistic insights into the APP-driven promotion of neurite outgrowth and neuronal differentiation, relevant for neuroregeneration.

## Supporting information

Online Resource

## Conflicts of interest/Competing interests and consent for publication

The authors have no conflict of interests to declare. All authors have approved the final version of the manuscript and give their consent to be published.

## Ethics approval

Animal-related ethics are included in the ‘Cortex dissection and APP-pro-EGF co-Immunoprecipitation’ sub-section of the Materials and Methods section.

## Authors’ contributions

JR performed the EGF experiments, LB built the MycHB-EGF construct and performed the HB-EGF assays, SD performed the YTH screens and helped on the yeast co-transformation assays, ARB performed the real time PCR assays, under SIV supervision, with the help of OAB, and of UK (rodents’ brains IPs). JR, LB, ARB and SIV analyzed and interpreted all data, and wrote the manuscript, which all authors have revised.

