## Supplementary material for "APP binds to the EGFR ligands HB-EGF and EGF, acting synergistically with EGF to promote ERK signaling and neuritogenesis": Online Resource

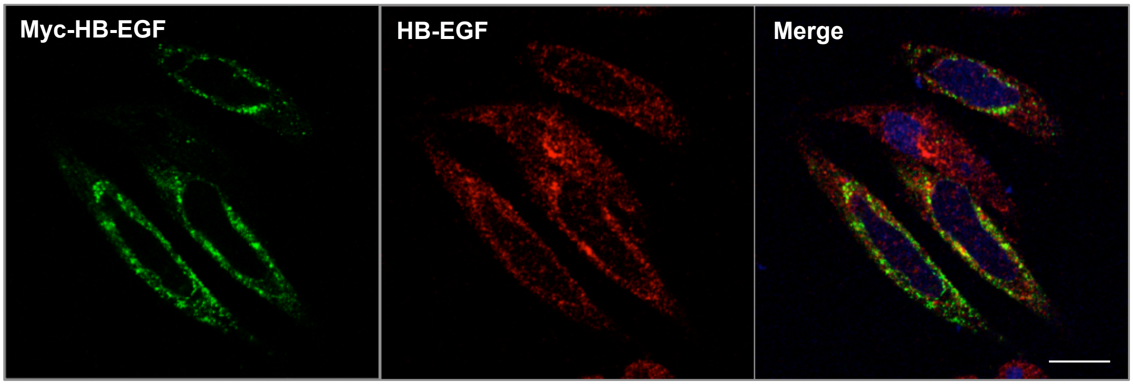


**Supplementary Figure S1. Subcellular distribution and co-localization of exogenous Myc-HB-EGF with endogenous HB-EGF.** HeLa cells transiently overexpressing the Myc-HB-EGF cDNA were immunolabeled with the anti-Myc tag antibody (green) and with the anti-HB-EGF antibody (red). Bar = 10 µm.

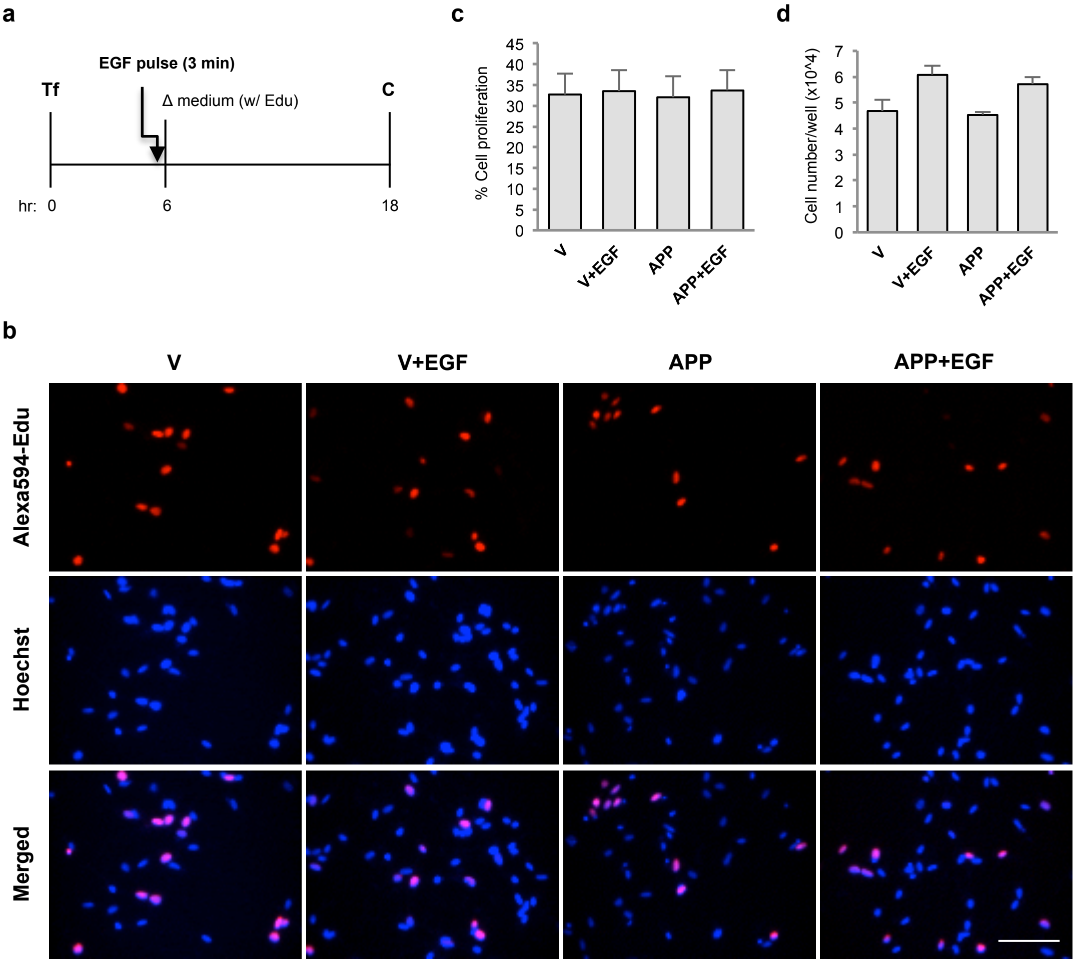


**Supplementary Figure S2. Effects of EGF treatment and APP overexpression on SH-SY5Y cell proliferation and survival. a** SH-SY5Y cells were either transfected ('Tf') with the GFP empty vector ('V') or APP695-GFP ('APP') cDNAs, and stimulated or not for 3 min with 100 ng/mL human recombinant EGF ('V+EGF' and 'APP+EGF') immediately before cell media change. Fresh media included the Click-iT® EdU, a nucleotide analogue incorporated in proliferating cells, and cells were fixed processed for analysis after 12h. **b** Representative epifluorescence microphotographs of cells tested with the Click-iT® EdU Alexa Fluor® imaging kit. Edu is labelled with a red fluorescing dye, and stains proliferating cells. Blue fluorescing Hoechst 33342 was used to assed the total number of cells. Bar = 50 µm. **c** Percentage of cell proliferation was calculated by counting the number of EdU red fluorescing cells versus the total number of Hoechst blue fluorescing cells (scored in 20 fields of view). **d** Plot of the total number of Hoechst positive cells per well in each experimental condition (scored in 20 fields of view). n = 3. Statistically non-significant results by the one-way ANOVA. Data is presented as mean ± standard error of the mean.

**a.** BetaIII-tubulin levels **b.** ERK1/2 phosphorylation

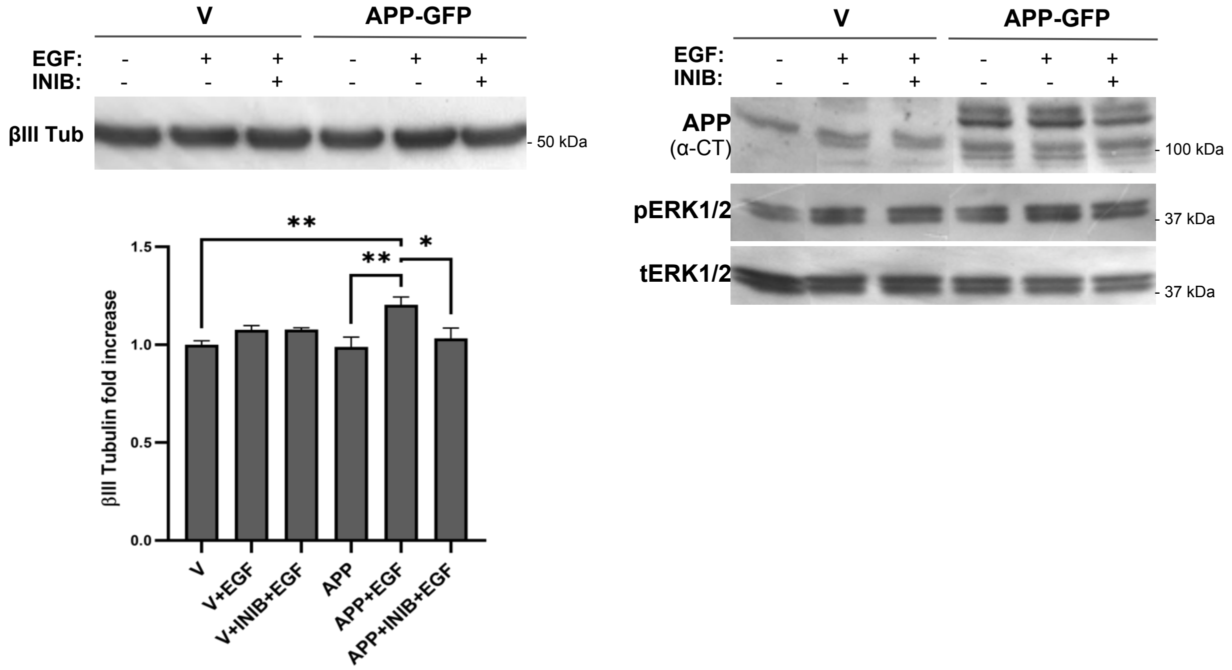


**Supplementary Figure S3. APP and EGF combined treatment increase SH-SY5Y neuronal-like differentiation via EGFR.** SH-SY5Y cells, either transfected with the GFP empty vector ('V') or APP695-GFP ('APP') cDNAs, were stimulated or not for 3 min with 100 ng/mL of human recombinant EGF ('V+EGF' and 'APP+EGF') immediately before cell media change (16hr before fixation/harvesting). When indicated, this EGF stimulation was performed under EGFR inhibition with 10 μM PD168393 ('INIB'), added 1h before EGF. Cells were collected for analysis after 24h of cell transfection. **a** βIII-tubulin protein levels under these conditions. Upper: immunoblots of cell lysates, probed with the βIII-tubulin antibody. Lower: βIII-tubulin protein levels were plotted as fold increases of the non-treated GFP-expressing cells ('V'). Of note, all the lanes are from the same blots, being rearranged into the present order. Ponceau S staining of total proteins bands was used as loading control for all the data, and data is presented as mean ± standard error of the mean. n = 4-5. *$p < 0.05$, **$p < 0.01$ by one-way ANOVA followed by the Tukey's multicomparison post-test. **b** Immunoblots of cell lysates, probed with the antibodies against APP C-Terminus and against phospho-ERK1/2 ('pERK') and total ERK 1/2 ('ERK') (graphic with quantitative determinations presented in Fig. 6). Of note, all the lanes are from the same blots, being rearranged into the present order.
